# The activation of fibroblast growth factor receptor 1 provides therapeutic benefit in a mouse model of Alzheimer’s disease

**DOI:** 10.1101/2025.02.17.638757

**Authors:** Wei-Chien Hung, Wei-Chun Sun, Tzu-Ching Su, Yi-Jung Lee, Irene Han-Juo Cheng

## Abstract

Alzheimer’s disease (AD) is a neurodegenerative disease that is characterized by the accumulation of amyloid plaque and neurofibrillary tangles, ultimately impairing multiple cognitive domains. Both plaques and tangles cause neuronal damage and stimulate inflammatory responses in glial cells. The fibroblast growth factor receptor 1 (FGFR1)-mediated signaling pathways support the function of damaged neurons and modulate inflammatory response. The FGFR1 agonists, including Fibroblast growth factor 1 (FGF1) and FG loop peptide (FGL), have been implicated in multiple disease therapy. Whether FGFR1 agonists can improve pathology and cognitive function in AD remains unknown. Here, we showed that administration of FGF1 and FGL to the AD mouse model reversed spatial memory impairment, enhanced neurogenesis, suppressed reactive astrogliosis, and restricted dystrophic neurite. However, only FGF1 treatment reduced the deposition of senile plaque. In microglial culture studies, FGF1 improves the phagocytosis ability of microglia, which contributes to the clearance of plaques. Together, our findings suggested that FGFR1 agonists alleviate pathology and cognitive impairment via immunomodulatory and improve neuronal health in the AD mouse model.

**Highlights:** FGFR1 agonists alleviate memory deficits in AD mice.

FGFR1 agonists increase adult neurogenesis and limit dystrophic neurites in AD mice.

FGFR1 agonists increase plaque-associated microglia *in vivo* and promote phagocytosis *in vitro*.

## 1. Introduction

Alzheimer’s disease (AD) is the most common form of dementia, with onset primarily occurring after the age of 65. Symptoms of AD include impaired cognitive functions, such as memory loss and language difficulties (“2024 Alzheimer’s disease facts and figures,” 2024). The neuropathological hallmarks and definite diagnostic criteria of AD are the depositions of extracellular senile plaques and intracellular neurofibrillary tau tangles. Senile plaques consist of aggregated amyloid-β protein (Aβ), which is derived from the cleavage of amyloid precursor protein (APP). Altered Aβ homeostasis leads to distinct AD progression. In the healthy brain, Aβ production and clearance are tightly balanced. In the AD brain, impaired Aβ clearance leads to its accumulation.

The abnormal accumulation of Aβ impairs neuronal function and reduces neurogenesis. These toxic amyloid plaques induce neurodegeneration, predominantly at axon terminals located near the plaques, term neurite plaques (Cheng et al., 2007; Mabrouk et al., 2023). These dystrophic neurites have swollen, bulbous-shaped neuronal processes containing neurofilament proteins, tau, APP, and lysosomal vesicles (Lie & Nixon, 2019). Furthermore, the pathological microenvironment created by Aβ disrupts neuronal maturation, thereby impairing neuronal proliferation and differentiation (Moon et al., 2014). Impaired adult neurogenesis is noticed in AD patients and correlates with cognitive decline (Salta et al., 2023). In adult mammals, neurogenesis remains active in the subgranular zone (SGZ) of the hippocampal dentate gyrus (DG). Adult neurogenesis in the SGZ significantly contributes to memory formation and the learning process (Anacker & Hen, 2017), which is impaired in the AD mouse model (Deng et al., 2022). Besides Aβ, cytokine and growth factors secreted from glial cells could also modulate adult neurogenesis (Araki et al., 2021), suggesting the interaction between glial cells and neurons in the hippocampus under AD progression.

The accumulation of Aβ plaques activates microglia and astrocytes to induce neuroinflammation, which is one of the key factors in the pathogenesis of AD (Cameron & Landreth, 2010; Patani et al., 2023). Microglia are the innate immune cells surveying the central nervous system (CNS) to clear accumulated Aβ through phagocytosis (Podlesny-Drabiniok et al., 2020). Numerous studies have suggested the protective role of microglia around plaques by limiting the spread of toxic amyloid (Huang et al., 2021; Liu et al., 2017). By forming the microglia barrier, plaque-associated microglia build the physical wall to restrain the Aβ accumulation (Condello et al., 2015). However, prolonged exposure to Aβ shifts microglia to a reactive status, resulting in reduced efficiency in Aβ clearance and increased production of pro-inflammatory cytokines (Fruhwurth et al., 2024). On the other hand, reactive astrocytes also respond to pathological stimuli by extending the hypertrophic process in the AD brain (Smit et al., 2021). Astrocytes have multifaceted functions in regulating metabolism, synaptogenesis, and neurogenesis (Han et al., 2021). Aβ accumulation triggers a specific morpho-functional remodeling of astrocytes—a process known as astrogliosis. Reactive astrocytes regulate Aβ deposition and inflammatory response in AD mouse model (Patani et al., 2023).

Fibroblast growth factor receptors (FGFR1-4) are receptor tyrosine kinases involved in multiple cellular processes (Xie et al., 2020). FGFR1-3 are widely expressed in the brain, while FGFR4 is absent in the adult brain. In the CNS, FGFR1 is widely expressed in microglia, astrocytes, and neurons, modulating the protective signaling pathways (Klimaschewski & Claus, 2021). In neuronal culture, activating FGFR1 promotes neurite outgrowth and neuronal survival (Pankratova et al., 2016; Saffell et al., 1997). Neuronal *Fgfr1* knockout mice are impaired in neurogenesis, long-term potentiation (LTP), and memory consolidation (Zhao et al., 2007). In astrocytes and microglia, FGFR1 controls their activation into neuroprotective phenotypes under stress (Zhou et al., 2025; Zou et al., 2019). One of the main ligands of FGFR1 in the brain is FGF1. Intraperitoneal-given FGF1 improves diabetes-induced cognitive decline (Wu et al., 2020). An artificially modified FGF1, named aFGF135, was developed to treat spinal cord injury in adult paraplegic rats and human patients (Ko et al., 2018; Wu et al., 2011). Because FGFR1 is downregulated in the hippocampus of AD patients (Ho Kim et al., 2015), we propose that FGFR1 agonists could offer a beneficial effect for AD. However, FGF1 has a short half-life *in vivo,* and it is difficult to penetrate the blood-brain barrier (BBB) (Zakrzewska et al., 2009). We thus also want to examine the efficacy of another FGFR1 agonist, FG loop peptide (FGL), which is a 15-amino acid peptide (EVYVVAENQQGKSKA) designed from the neural cell adhesion molecule (NCAM) (Neiiendam et al., 2004). FGL is stable and can penetrate the BBB to increase FGFR1 phosphorylation (Secher et al., 2006). In multiple rodent models, FGL injection promotes synaptogenesis, enhances LTP, facilitates memory, and reduces neuroinflammation (Cambon et al., 2004; Downer et al., 2010; Knafo et al., 2012; Pereda-Perez et al., 2019; Popov et al., 2008; Secher et al., 2006).

In the present study, we stimulated the FGFR1 with FGF1 or FGL through different routes in the AD mouse model. Hippocampal injection of FGF1 improves memory and pathological deficits of APP transgenic mice. Subcutaneous injection of FGL yielded partial improvements in these deficits. We demonstrated that stimulating the FGFR1 pathway might be a potential therapeutic approach for AD.

## 2. Materials and methods

### 2.1. Animals

APP transgenic mouse line J20 (B6.Cg-Zbtb20^Tg(PDGFB-APPSwInd)20Lms^/2Mmjax), carrying the human APP gene with the Swedish (K670N/M671L) and India (V717F) familial mutations, was used as an AD mouse model. Mice were kept under 12-hour light-dark cycle at 25°C and 60% humidity, with ad libitum access to food and water. For four months of age groups, FGF1 was given by stereotactic injection into the hippocampus of mice. Morris water maze was carried out 1 month after the FGF1 administration. For seven months of age groups, FGL was given by subcutaneous injection into the mice every other day for 23 days, including the behavior test period. One day after the behavior test, mice were anesthetized by intraperitoneal urethane injection and perfused with normal saline for 5 minutes. Brain tissues were collected for immunohistochemistry staining and ELISA. The study was approved by the Institutional Animal Care and Use Committee (IACUC) of National Yang Ming Chiao Tung University. All experimental procedures involving animals and their care were carried out following the Guide for the Care and Use of Laboratory Animals published by the United States National Institute of Health (NIH).

### 2.2. FGF1, FGL, and control peptide preparation

Human FGF1, a single nonglycosylated polypeptide chain with 135 amino acids, was prepared by the company EUSOL as previously described (Ko et al., 2018). FGL peptide (EVYVVAENQQGKSKA) and control peptide (FGL-ala, EVYVVAENAAGKSKA) were dissolved in distilled water (5 mg/ml). The FGL-ala design substitutes double alanine for double glycine on the 9^th^ and 10^th^ amino acids, the critical site for interacting with FGFR (Kiselyov et al., 2003). FGL-ala has been proven to have no effects on neurite growth (Kiselyov et al., 2003), synaptic function, cognitive function (Cambon et al., 2004), and Aβ-induced pathology (Klementiev et al., 2007). FGL-ala has been commonly used as a negative control of FGL in various studies. The injected form of FGL peptide comprises two FGL monomers coupled to Aminodiacetic acid (N-(carboxymethyl) glycine) through their N-terminal (Neiiendam et al., 2004) by Mission Biotech, Taiwan.

### 2.3. Stereotactic surgery

4-month-old mice were maintained under single housing conditions before surgery. Mice were anesthetized with isoflurane and placed into a stereotaxic frame. The small holes were drilled at +/− 1.3 mm ML and −2.0 mm AP from bregma; −1.5 mm DV from the brain surface. FGF1 (3 μg) or vehicle (0.5 μL) were injected into CA1 of mice (0.5 μl, bilateral) with a Hamilton microsyringe at a rate of 0.5 μl/minute. After injection, the needle was retained on the injection site for 10 minutes and slowly withdrawn at 0.5 mm/minute. Closing the skin incision using nylon sutures and monitoring the mouse condition 24 hours after surgery. Behavior experiments were conducted 1 month following surgery.

### 2.4. Morris water maze

The water maze contains a circular pool (diameter 120 cm) filled with water. A hidden platform (diameter 10 cm) was submerged 1 cm below the water level in the center of the target quadrant. On days 1-5, mice went through two sessions per day and three training trials per session. In each trial, mice were allowed to search for the platform for 1 minute and forced to stay on the platform for 15 seconds. If mice could not find the platform within 1 minute, they were guided to it. On day 6, the platform was removed, and mice were put into the pool in the opposite quadrant of the target quadrant and allowed to swim for 1 minute. On days 7-9, the platform was placed in the opposite quadrant. Mice went through two sessions per day and three training trials per session. The tracks and the time were recorded and analyzed by Ethovision software (Noldus, Wageningen, the Netherlands).

### 2.5. Cell culture

BV2 microglial cell line (Cat. No ABC-TC212S, AcceGen Biotech, USA) was maintained in RPMI supplemented with 10% FBS, 1% L-glutamine (GLL01, Caisson laboratory, USA), and 3.7g sodium bicarbonate. Cell lines were incubated at 37°C in a humidified 5% CO2 atmosphere.

### 2.6. Primary microglia culture

Primary microglia were cultured from C57B6/J mice with postnatal days 0-3. The cortex was digested with 100U papain and 400U DNase I in Hanks’ Balanced Salt Solution (HBSS) buffer at 37°C for 30 minutes. Digested cells were passed through a 70-μm cell strainer (Corning, USA). Cortical mixed cells were grown in DMEM/F12. After 21 days of incubation, microglia were isolated by 5% trypsin/ Phosphate-buffered saline (PBS) and were seeded in 4-well plates for 24 hours.

### 2.7. Oligomeric Aβ (oAβ) preparation

Hexafluoroisopropanol (HFIP)-treated Aβ1-42 peptides (A-1163-2, rPeptide Inc, USA) were dissolved in 10% Dimethyl sulfoxide (DMSO) at 100 μM and stored at –80°C. Before the experiment, the stock was put at 4°C for 24 hours to generate oAβ.

### 2.8. Western blotting

The hippocampus and cortex were dissected from the frozen brain and homogenized in RIPA buffer with phosphatase and protease inhibitors (Roche). Protein was separated by 8% Tris-glycine SDS-polyacrylamide gel and transferred to the nitrocellulose membranes (Nitropure, MSI Corp., USA). After blocking in casein buffer for 2 hours, anti-FGFR1 antibody (9740, Cell signaling, USA) and anti-β-actin antibody (2D4H5, 66009-1-Ig, proteintech, USA) was incubated with the membranes at 4°C overnight. The membranes were washed by TBST three times and incubated with the HRP-conjugated secondary antibodies for 1 hour. The signal was developed by the chemiluminescent (ECL) substrate (GTX14698, GeneTex, USA) and detected by a luminescence imaging system (LAS-4000, Fujifilm, Japan).

### 2.9. Enzyme-linked immunosorbent assay (ELISA)

Frozen hippocampal samples were homogenized in 5 M guanidine/5 mM Tris buffer (pH 8.0). The samples were diluted with 0.25% casein blocking buffer containing 0.5 M guanidine and protease inhibitor (04693116001; Roche, Switzerland) to 50X working concentration. Total Aβ and Aβ 1-42 levels were detected by ELISA kits (27729 and 27711; IBL, Japan) according to the manufacturer’s instructions.

### 2.10. Immunohistochemistry

Brain tissues were fixed in 4% paraformaldehyde overnight, immersed in 15% sucrose for 24 hours, and then changed to 30% sucrose. All the sections were sliced coronally at a thickness of 30 μm on a sliding microtome. Subsequently, brain slices were selected and blocked with PBS containing 10% FBS (1210466; Thermo Fisher Scientific Gibco, MA, USA) and 0.5% triton X-100 for 1 hour, then overnight incubation at 4°C with primary antibody, followed by the incubation with secondary antibody for 1 hour. Primary antibodies used were: anti-Iba1 (019-19741, Wako, Japan) antibody, anti-GFAP antibody (Z0334, Dako, USA; 3670, Cell signaling, USA), anti-LAMP1 (1D4B, DSHB, USA), anti-Aβ (D54D2: 8243, Cell signaling, USA; MOAB2: ab126649, Abcam, USA), and anti-DCX antibody (sc-8066, Santa Cruz Biotechnology, USA; 4604, Cell signaling, USA), Secondary antibodies used were: Alexa Fluor 488 goat anti-rabbit IgG, Alexa Fluor 594 goat anti-rabbit IgG, Alexa Fluor 488 goat anti-mouse IgG, Alexa Fluor 594 goat anti-mouse IgG, Alexa Fluor 405 goat anti-mouse IgG, Alexa Fluor 594 goat anti-rat IgG, and Alexa Fluor 488-conjugated AffiniPure Donkey Anti-goat IgG. The slices were mounted and visualized using fluorescence microscopes (Axio Observer A1; Zeiss, Germany, BX63; Olympus, Japan). Microglia coverage, dystrophic neurite, and neurogenesis were quantified by ImageJ software (NIH). Microglia coverage was calculated by the Iba1-positive cell within a circular area with a radius of 25 μm centered on plaques. Dystrophic neurite was identified by the percentage of LAMP1-positive area within a circular area with a radius of 30 μm centered on plaque normalized to per Aβ plaque area in the hippocampus. Neurogenesis was determined by the number of DCX-positive cells in the DG per slice.

### 2.11. Microglia phagocytosis assay and immunocytochemistry

Purified primary microglia or BV2 cells were seeded at a density of 10^5^ cells/well on poly-D-lysine coated coverslips at 4-well plates. The attached cells were treated with a control buffer or FGF1 for 24 hours. The phagocytic ability of each cell type was examined by incubating with fluorescent carboxylated microspheres (F8821, 1 μm in diameter, Invitrogen, USA), which pre-coated with FBS at 37°C for 30 minutes. For fluorescence-conjugated Aβ phagocytosis, BV2 cells were seeded on poly-D-lysine coverslips in a 4-well plate overnight. After the cells were attached, the medium was replaced with a medium containing 5 μM oAβ and incubated for 24 hours. The Aβ phagocytosis was examined by incubating the FGF1-treated cells with 0.2 μM fluorescence-conjugated Aβ (AS-60479-01, ANASPEC, USA) for another 24 hours. After three times of PBS washes, attached cells were fixed with 4% paraformaldehyde. Cells were stained with anti-Iba1 antibody overnight at 4°C, followed by the incubation with Alexa Fluor 594 goat anti-rabbit IgG secondary antibody for 1 hour or phalloidin (#00045, Biotium, USA) for 20 minutes to visualize the number of engulfed microspheres in the cell.

### 2.12. Statistical analysis

GraphPad Prism (version 8.0; GraphPad, USA) was used for statistical analysis. Differences among multiple means were assessed by one-way, two-way ANOVA. Differences between the two means were assessed by an unpaired t-test. All data are presented as mean ± SEM. The threshold for significance was defined as *p* < 0.05.

## 3. Results

### 3.1. The expression of FGFR1 decreased in the hippocampus of APP mice

The expression of FGFR1 has been reported in both mice and humans. We extracted the RNAseq results from the HPA mouse dataset (https://www.proteinatlas.org/). Of all brain regions, FGFR1 has the highest expression in the hippocampus (Figure 1a), which is the critical region for memory formation and most particularly vulnerable in AD. Because FGFR1 is down-regulated in the hippocampus of AD patients (Ho Kim et al., 2015), we examined the change of FGFR1 in the human APP transgenic mice model (line J20). Immunoblot analysis revealed significantly decreased FGFR1 in the hippocampus of APP mice compared to wildtype (WT) littermate controls at both seven (*p* = 0.0332, Figure 1b, c) and ten months old (*p* = 0.0273, Figure 1b, d). Given the beneficial effect of FGFR1 signaling, we hypothesized that stimulating FGFR1 may ameliorate functional and pathological deficits of APP mice.

**Figure 1.**
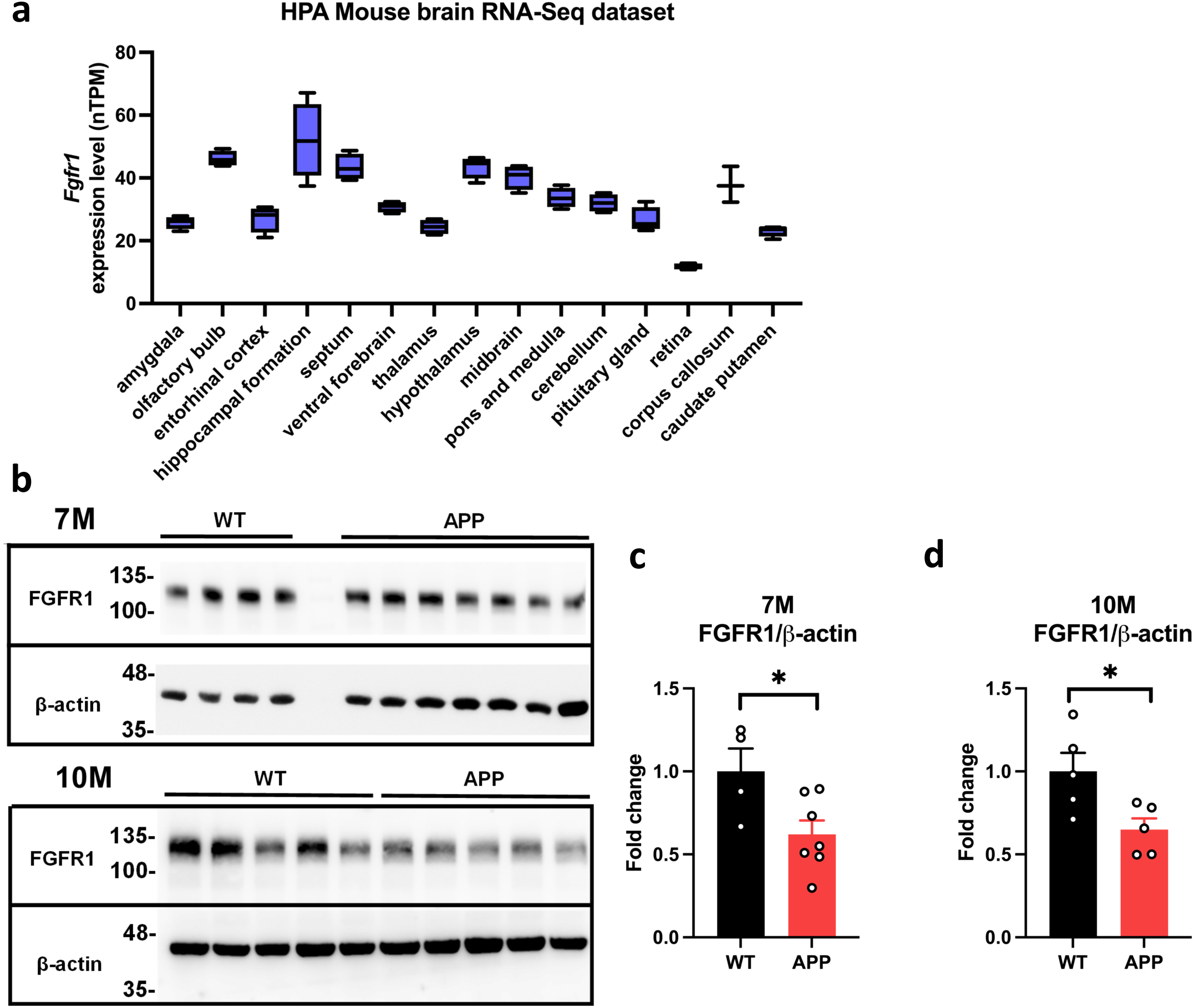
FGFR1 is highly expressed in the hippocampus and reduced in the hippocampus of APP mice. (a) *Fgfr1* levels in different brain regions extracted from HPA mouse brain RNA-Seq dataset. (b) Representative Western blotting images of the FGFR1 in the hippocampus of 7 and 10-month-old WT or APP mice. Actin was used as the loading control. (c-d) Quantification of FGFR1 levels in the hippocampus of 7 (c) and 10 (d) -month-old WT or APP mice. N = 4-7 per group. Results were analyzed by unpaired *t*-test. * *p* <0.05.

### 3.2. FGF1 administration rescued spatial memory deficit and Aβ accumulation

A previous study revealed that intraperitoneal-given FGF1 improves spatial memory in diabetic mice (Wu et al., 2020). We checked whether applying FGF1 in hippocampal parenchyma can reverse the memory deficiency in the AD mouse model. WT and APP mice were microinjected with a single dose (3 μg) of FGF1 or saline control into the CA1 region of the hippocampus at 4 months old. Morris water maze (MWM) tests were conducted to examine whether the FGF1 administrate could rescue the memory deficits of AD mice at 5 months old and sacrificed for biochemical and histopathological analysis at 5.5 months old (Figure 2a). APP mice had memory deficiency in the acquisition phase compared to WT mice. FGF1-injected APP mice restored this deficiency on the 4^th^ and 5^th^ day of training (*p* = 0.0181 and 0.016, respectively, Figure 2b). In the probe test on day 6, the latency to the platform is significantly longer in APP mice than in WT mice (*p* < 0.0001, Figure 2c). FGF1 injection reversed this spatial memory deficiency in APP mice (*p* = 0.0429, Figure 2c).

**Figure 2.**
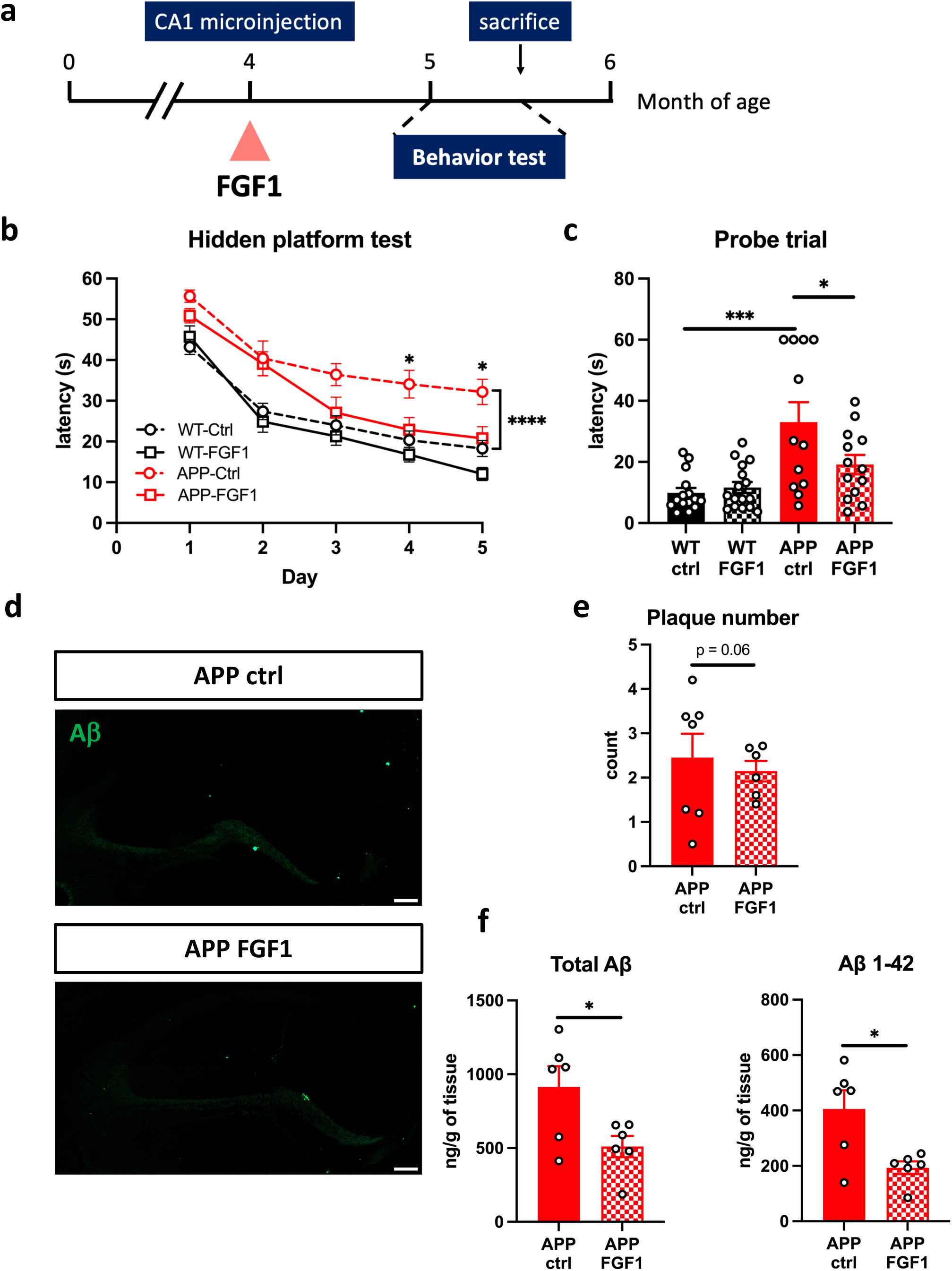
Hippocampal injection of FGF1 alleviated cognitive-related deficits in APP mice. (a) Timeline for FGF1 injection experiments. (b) The learning curves of the 5-day MWM hidden platform test. Results were analyzed by Two-Way ANOVA with Tukey post-hoc analysis. * *p* <0.05; **** *p* <0.0001. (c) The latency to the platform in the day 6 probe trial. N = 12-16 mice per group. Results were analyzed by One-Way ANOVA. * *p* <0.05; *** *p* <0.001. (d) The representative immunofluorescence images of Aβ plaque in the hippocampus. Scale bar = 200 μm. (e) The average number of plaques in the hippocampus per brain slice. N = 5-6 slices per mouse, 6-7 mice per group. Results were analyzed by unpaired *t*-test. (f) Levels of total Aβ and Aβ 1-42 in the hippocampus were determined by ELISA. N = 6 mice per group. Results were analyzed by unpaired *t*-test. * *p* <0.05.

Immunohistochemistry was applied to evaluate if amyloid plaque pathology was altered by FGF1 injection in APP mice. We observed a non-significant trend toward decreased amyloid plaque counts in the hippocampus of APP mice (*p* = 0.0613, Figure 2d, e). To measure the level of different Aβ species, ELISA was employed to determine the levels of guanidine-soluble total Aβ and Aβ42. Notably, the levels of total Aβ (*p* = 0.0282) and Aβ42 (*p* = 0.0134) were significantly reduced by FGF1 injection in APP mice (Figure 2f). These results suggest that FGF1 ameliorated cognitive functions and Aβ pathology in APP mice.

### 3.3. FGF1 reversed pathological deficits on multiple cell types in APP mice

Aβ induces pathological changes in multiple cell types of AD patients and APP mice models, including plaque-associated gliosis and neuronal abnormality. Reactive astrocytes appeared to increase in the late stage of AD patients (Kumar et al., 2023). We examined the activation of astrocytes in the hippocampus. Brain slices from all four groups of mice were immune stained using antibody against GFAP, a commonly used marker for astrocytes. An increase in GFAP expression was observed in multiple neurodegenerative diseases (Simpson et al., 2010). In the control groups, reactive astrogliosis coverage was significantly higher in APP mice than in WT mice (*p* < 0.0001, Figure 3a, b). FGF1-injected APP mice had significantly lower reactive astrogliosis coverage than APP mice (*p* = 0.0194, Figure 3a, b). To understand the effect of FGF1 on microglia-plaque interaction, we checked the microglial coverage surrounding the plaques. APP mice injected with FGF1 had higher microglia coverage around the plaque than the control group (*p* = 0.0115, Figure 3c, d). These results indicated that FGF1 decreases astrogliosis and triggers more microglia migrating toward the plaque.

**Figure 3.**
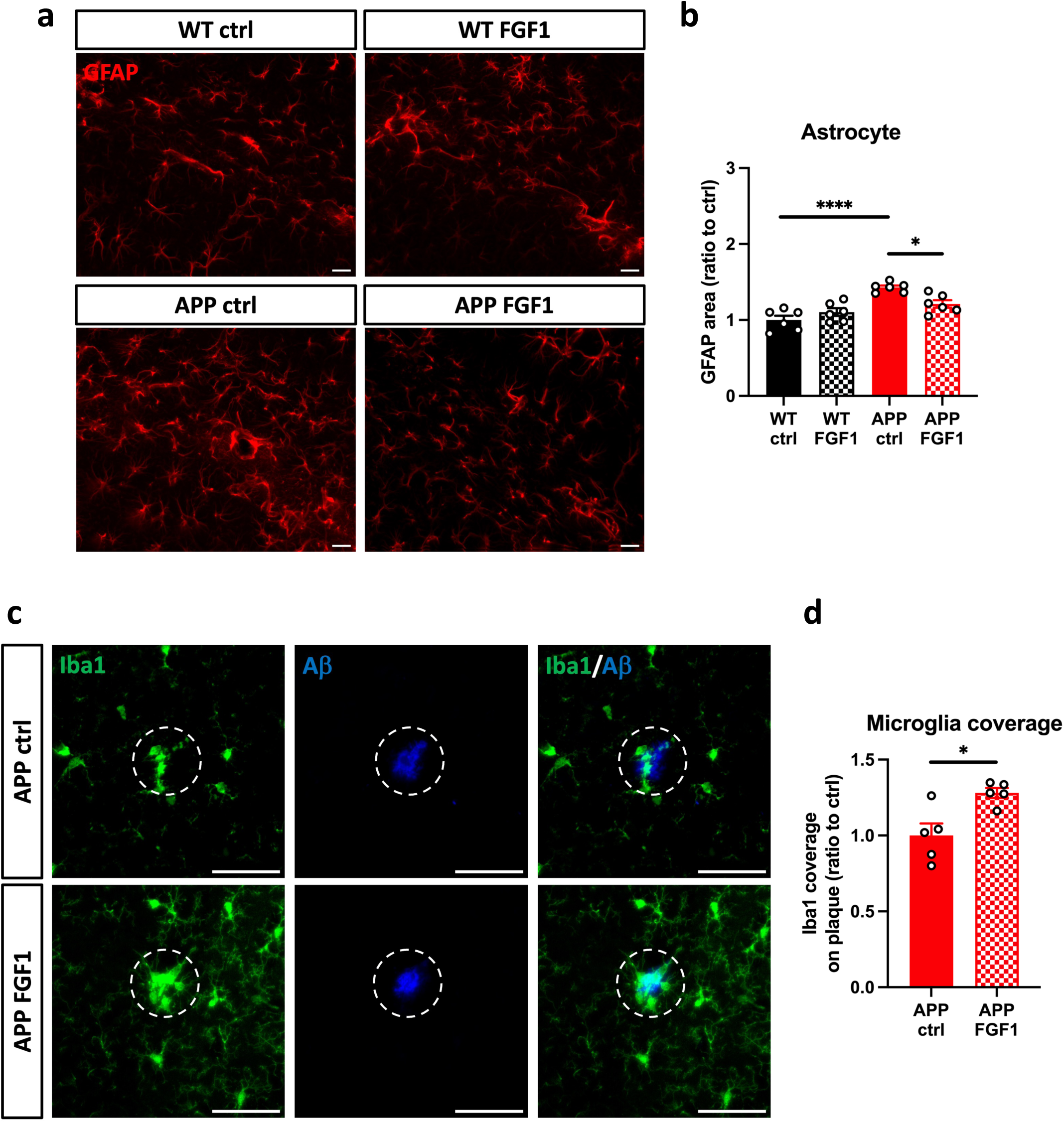
Hippocampal injection of FGF1 inhibited astrocyte activation and promoted plaque-associated microglia. (a) The representative immunofluorescence images of astrocytes in the hippocampus. Scale bar = 10 μm. (b) Quantification of the astrocytes area in the hippocampus. The normalized astrocyte coverage in the WT-ctrl group was set as 1. N = 6-10 slices per mouse, 6 mice per group. Results were analyzed by One-Way ANOVA. * *p* <0.05; **** *p* <0.0001. (c) The representative immunofluorescence images of microglia (green) coverage around the plaque (blue) in the hippocampus of APP-control mice and APP-FGF1 mice. Scale bar = 50 μm. (d) The area covered by microglia within a radius of 25 μm centered around the plaque. The average Iba1 positive area of APP-ctrl was set as 1. N = 7-9 plaques per mouse, 5 mice per group. Results were analyzed by unpaired *t*-test. * *p* <0.05.

Improved performance in the MWM test is commonly associated with the increase of neurogenesis in the DG of the hippocampus (Anacker & Hen, 2017), which could be regulated by FGFR1-mediated signaling (Kang & Hebert, 2015; Zhao et al., 2007). Thus, the brain sections were stained with DCX antibody to determine neurogenesis. In the control injected group, the number of DCX-positive cells in the DG was significantly lower in APP mice than in WT mice (*p* = 0.0013, Figure 4a, b), which indicated the reduced neurogenesis in APP mice. FGF1-injected groups induced more DCX-positive neurons than the control groups in both WT (*p* = 0.0407) and APP (*p* = 0.047) mice (Figure 4a, b). In addition, we evaluate the dystrophic neurites with LAMP1 antibody, a lysosome marker located at the lysosomal membrane. In APP mice, the LAMP1-immunoreactivity area surrounding the plaque was significantly lower in FGF1-injected mice than in the control injection group (*p* = 0.0357, Figure 4c, d). Together, FGF1 increases neurogenesis and restricts the spreading of neurite damage in APP mice.

**Figure 4.**
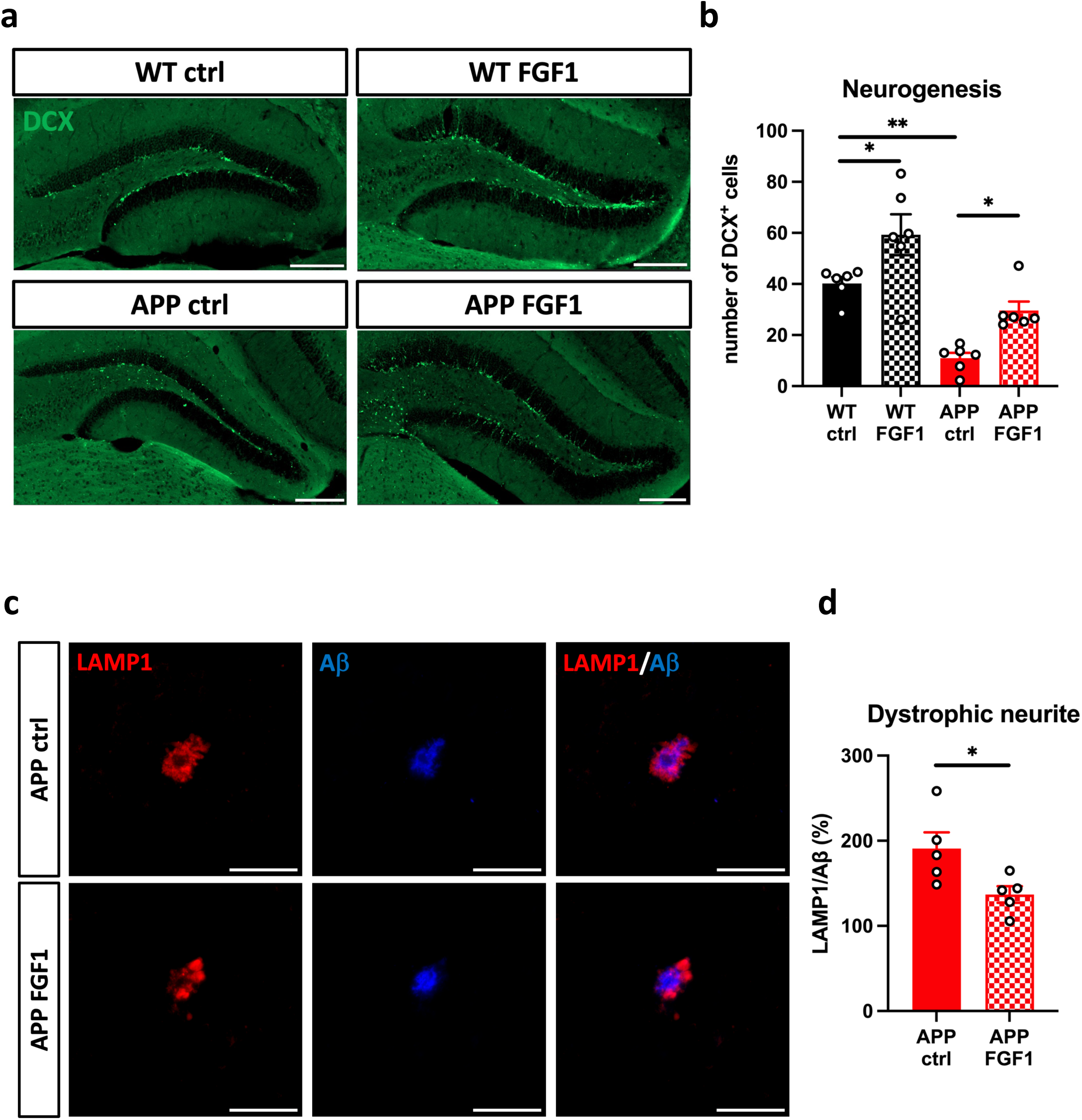
Hippocampal injection of FGF1 increased adult neurogenesis and limited dystrophic neurites spreading. (a) The representative immunofluorescence images of neurogenesis (DCX) in the DG. Scale bar = 200 μm. (b) Quantification of the number of DCX-positive cells per slice in the DG. N = 7-8 slices per mouse, 6 mice per group. Results were analyzed by One-Way ANOVA. * *p* <0.05; ** *p* <0.01. (c) The representative immunofluorescence images of dystrophic neurites. Scale bar = 50 μm. (d) Quantification of the percentage of plaque-surrounding dystrophic neurites in the hippocampus. The dystrophic neurite of APP-ctrl was set as 100%. N = 7-9 plaques per mouse, 5 mice per group. Results were analyzed by unpaired *t*-test. * *p* <0.05.

### 3.4. FGF1 enhances the phagocytic ability of microglia *in vitro*

The increase of plaque-associated microglia and decrease of Aβ deposition in FGF1-treated APP mice suggests that FGF1 may enhance phagocytosis of microglia to engulf amyloid plaques. Thus, primary microglial and BV2 cells were used to examine the impact of FGF1 on the phagocytosis ability *in vitro*. Microglial cells were treated with FGF1 for 24 hours and then incubated with fluorescence-labeled latex beads to detect the phagocytosis ability. FGF1-treated BV2 cells had a significantly higher percentage of cells ingested beads (*p* < 0.0001, Figure 5a, b) and more average number of beads per phagocytic cell (*p* = 0.0001, Figure 5a, c). Similarly, FGF1-treated primary microglia had a higher percentage of cells ingested beads (*p* = 0.0173, Figure 5d, e) and more average number of beads per phagocytic cell (*p* = 0.0336, Figure 5d, f) than control-treated cells. To specifically examine the Aβ phagocytosis, BV2 microglial cells were treated with FGF1 for 24 hours and then incubated with fluorescence-conjugated Aβ for 24 hours. FGF1-treated cells had significantly higher Aβ-positive area per ingested cell than control-treated cells, but no difference in the percentage of cells ingested Aβ (Figure 5g-i). These results indicated that FGF1 could improve the phagocytic ability of microglia to remove Aβ.

**Figure 5.**
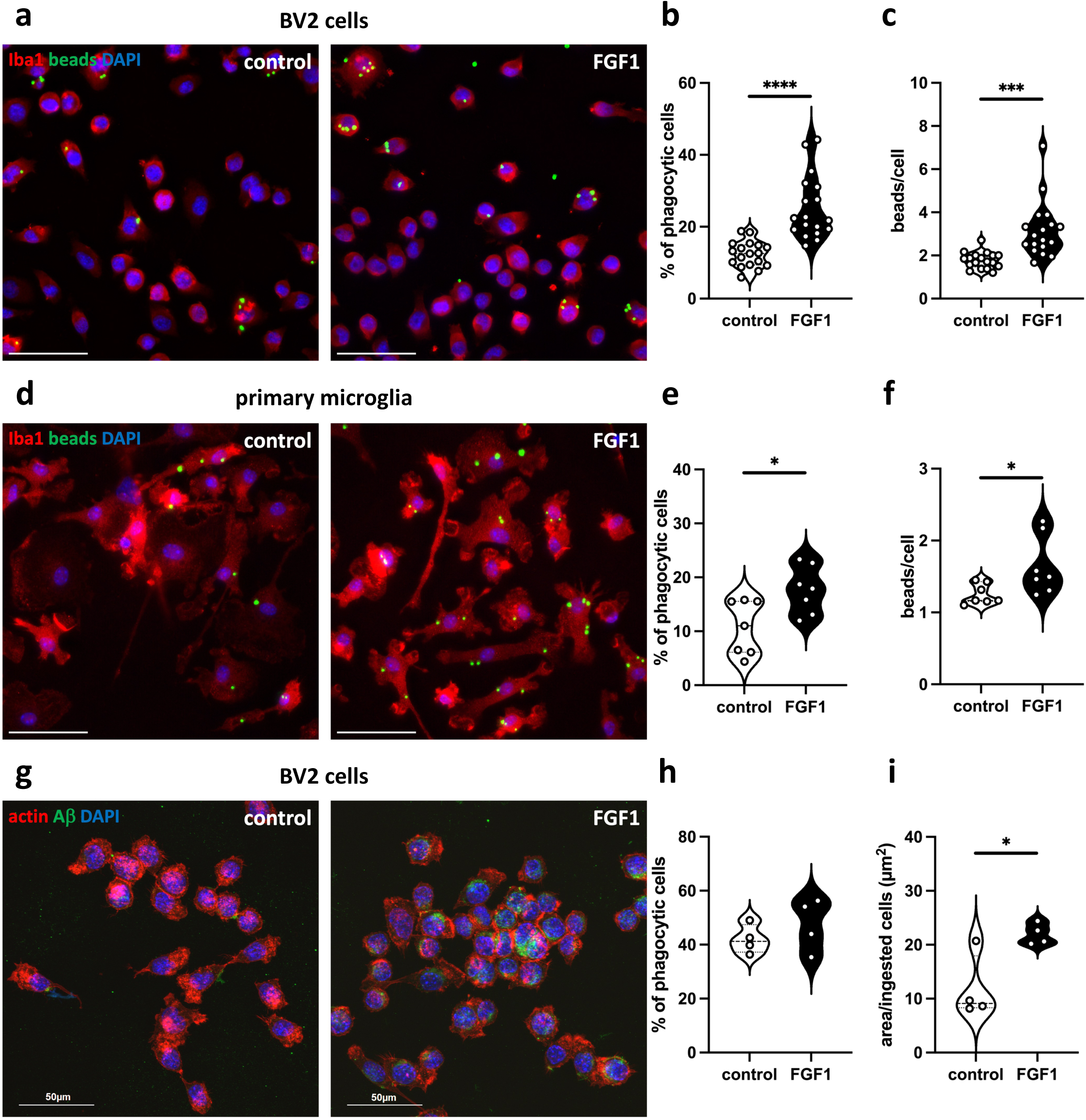
FGF1 increases phagocytosis in the microglial culture. (a, d) Representative images visualizing the phagocytosis of the fluorescence-labeled beads using BV2 microglial cell line (a) or primary microglia (d) treated with control buffer or FGF1. (b, e) FGF1 increases the percentage of cells engulfing beads. (c, f) FGF1 increases the number of beads per microglia. N = 18 wells per treatment from 3 independent experiments in BV2 cells; N = 7 wells per treatment from 2 independent experiments in primary microglia. (g) Representative images visualizing the phagocytosis of the fluorescence-labeled Aβ in BV2 cells treated with control buffer or FGF1. (h) FGF1 did not alter the percentage of cells engulfing Aβ. (i) FGF1 increases the Aβ area in the phagocytose cells. Scale bar = 50 μm. N = 4 wells per treatment from 2–3 independent experiments. Results were analyzed by unpaired *t*-test. * *p* <0.05; *** *p* <0.001; **** *p* <0.0001.

### 3.5. FGL ameliorates memory acquisition but not Aβ deposition

Due to the limitation of FGF1 to pass through BBB, peripheral administration of FGL, an FGFR1 agonist that can penetrate the BBB, was explored to expand therapeutic potential. We subcutaneously injected FGL (10 mg/kg) or FGL-ala control peptides (10 mg/kg) into 7-month-old WT and APP mice every other day for 23 days. These mice were subjected to the MWM with reversal learning test, biochemical, and histopathological analysis (Figure 6a). In the first 5-day hidden platform test, APP mice spent a longer time reaching the platform than WT mice. Notably, FGL-injected APP mice reversed acquisition deficits only on day 4 (day 4, *p* = 0.0084, Figure 6b). However, there is no significant difference among all groups of mice in the probe trial test on day 6 (Figure 6c). We further challenged these mice using a reversal task to assess their flexibility in adapting to new information. In the day 7-9 reversal learning, APP mice spent more time finding the platform than WT mice (*p* < 0.0001, Figure 6b). FGL injection reversed memory impairment at day 9 in APP mice compared to the control group (*p* = 0.0234, Figure 6b). These results indicated that peripheral administration of FGL improves the acquisition and flexibility of spatial memory in APP mice.

**Figure 6.**
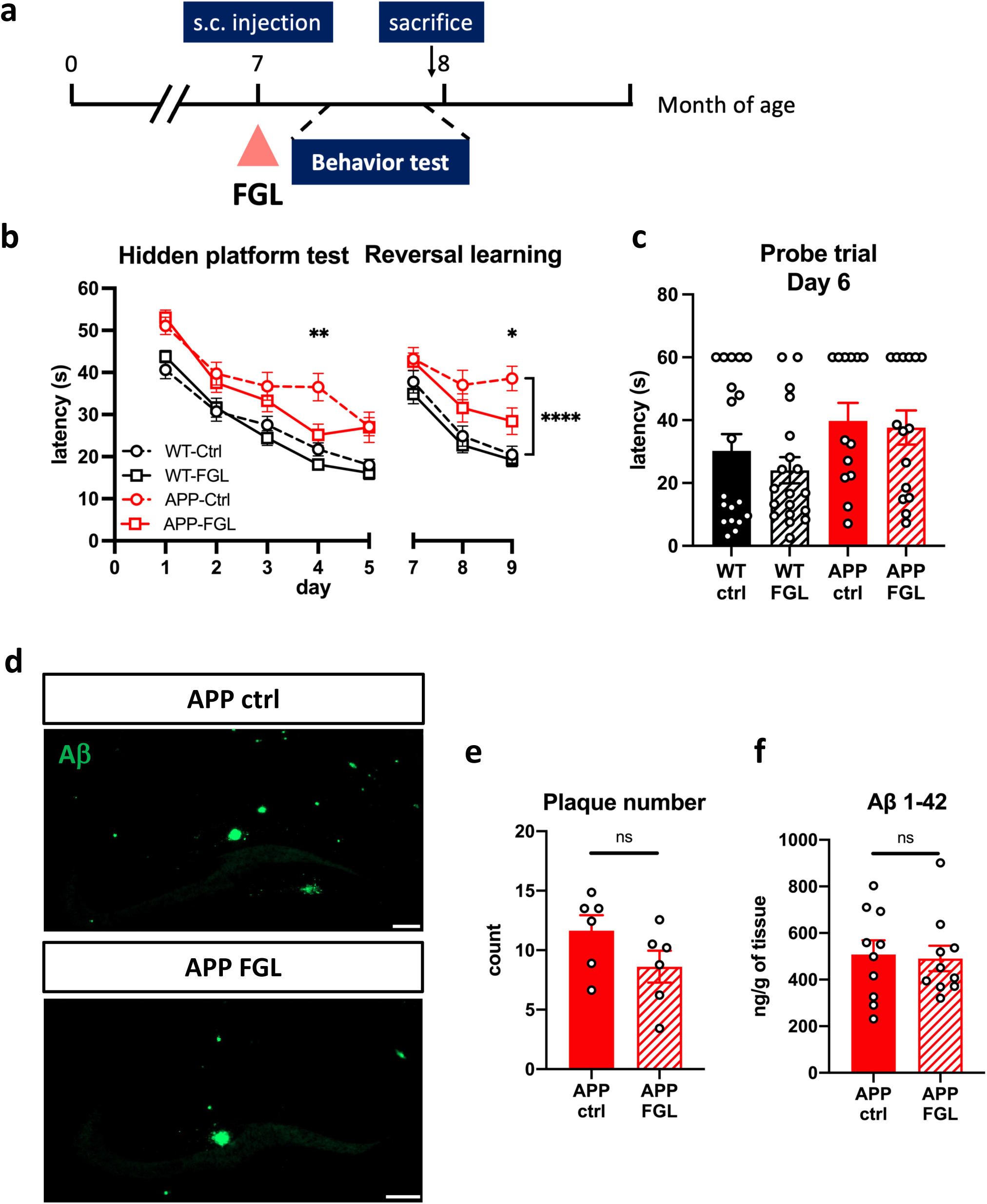
Subcutaneous injection of FGL partially alleviated cognitive-related deficits in APP mice. (a) Timeline for FGL injection experiments. (b) The learning curves of 5-day MWM training and 3-day reversal MWM training. Results were analyzed by Two-Way ANOVA with Tukey post-hoc analysis. * *p* <0.05; ** *p* <0.01; **** *p* <0.0001. (c) The latency to the platform in the day 6 probe trial. N = 13-19 mice per group. Results were analyzed by One-Way ANOVA. (d) The representative immunofluorescence images of Aβ plaque in the hippocampus. Scale bar = 200 μm. (e) The average number of plaques in the hippocampus per brain slice. N = 5-6 slices per mouse, 6 mice per group. (f) Levels of Aβ 1-42 in the hippocampus were determined by ELISA. N = 10 per group. Results were analyzed by unpaired *t*-test.

Next, we identified the influence of FGL on amyloid pathology. In contrast to the observed effects of FGF1, peripheral administration of FGL did not alter the amyloid plaque number (Figure 6d, e) in immunohistochemistry analysis, nor did it reduce the Aβ42 level in the ELISA assay (Figure 6f).

### 3.6. FGL reverses deficits on multiple cell types in APP mice

To compare with the FGF1 effect, we investigated whether peripheral administration of FGL impacts glial cells and neurons in APP mice. Similar to FGF1 administration, the FGL-injected APP mice had lower GFAP-positive coverage than control peptide-injected APP mice (*p* = 0.0418, Figure 7a, b), indicating the reduction of astrogliosis. For plaque-associated microglia, FGL-injected APP mice had significantly higher coverage of microglia surrounding the amyloid plaque than the control peptide-injected APP mice (*p* = 0.0054, Figure 7c, d), indicating the increase of microglia barrier formation.

**Figure 7.**
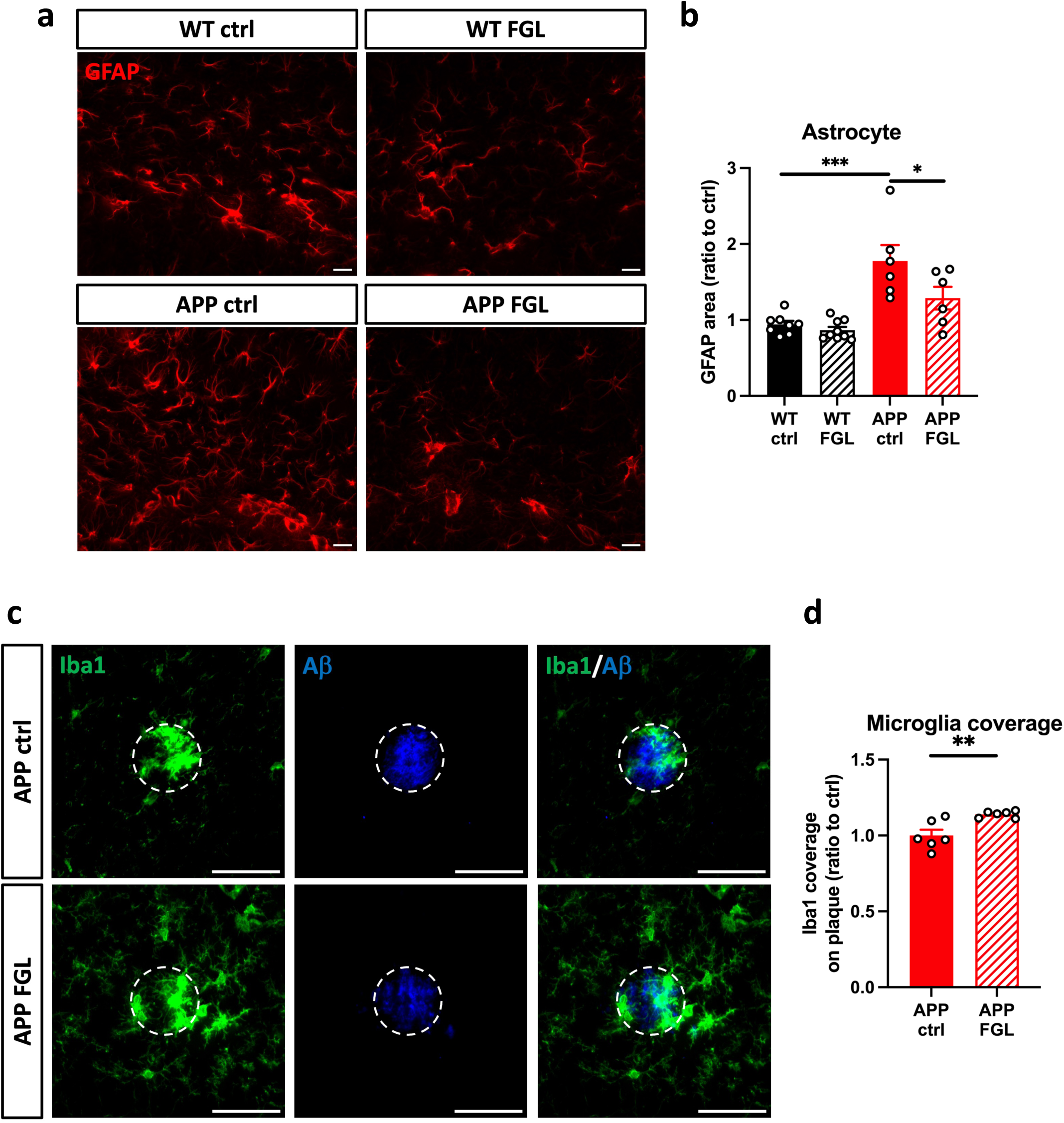
Subcutaneous injection of FGL inhibited astrocyte activation and promoted plaque-associated microglia. (a) The representative immunofluorescence images of astrocytes in the hippocampus. Scale bar = 10 μm. (b) Quantification of the area covered by astrocytes in the hippocampus. The normalized astrocyte coverage in the WT-ctrl group was set as 1. N = 5-7 slices per mouse, 6-9 mice per group. Results were analyzed by One-Way ANOVA. * *p* <0.05; *** *p* <0.001. (c) The representative immunofluorescence images of microglia (green) coverage around the plaque (blue) in the hippocampus of APP-control mice and APP-FGL mice. Scale bar = 50 μm. (d) The area covered by microglia within a radius of 25 μm centered around the plaque. The average Iba1 positive area of APP-ctrl was set as 1. N = 9-14 plaques per mouse, 6 mice per group. Results were analyzed by unpaired *t*-test. ** *p* <0.01.

To monitor neurogenesis, DCX-positive neurons in the DG were evaluated. The control-injected APP mice had significantly fewer DCX-positive neurons than the control-injected WT mice. FGL-injected APP mice had more DCX-positive neurons than control-injected APP mice (Figure 8a, b). In contrast to FGF1 injection, neurogenesis in WT mice was not altered by FGLs.

**Figure 8.**
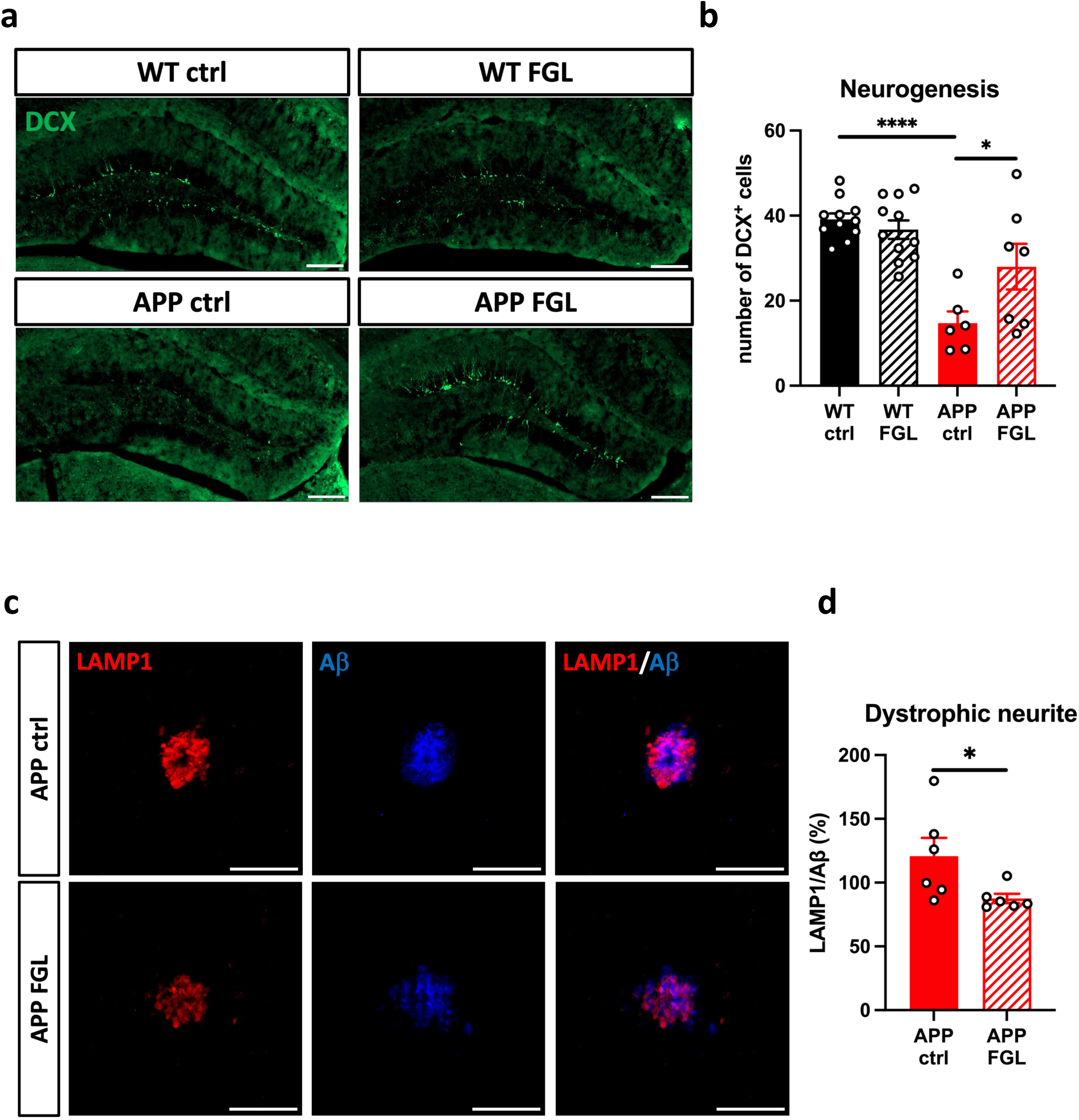
Subcutaneous injection of FGL increased adult neurogenesis and limited dystrophic neurites. (a) The representative immunofluorescence images of neurogenesis (DCX) in the DG. Scale bar = 200 μm. (b) Quantification of the number of DCX-positive cells per slice in the DG. N = 7-9 slices per mouse, 6-11 mice per group. Results were analyzed by One-Way ANOVA. * *p* <0.05; **** *p* <0.0001. (c) The representative immunofluorescence images of dystrophic neurites. Scale bar = 50 μm. (d) Quantification of the percentage of plaque-surrounding dystrophic neurites in the hippocampus. The dystrophic neurite of APP-ctrl was set as 100%. N = 9-14 plaques per mouse, 6 mice per group. Results were analyzed by unpaired *t*-test. * *p* <0.05.

For plaque-induced dystrophic neurites, we evaluated the LAMP1-immunoreactivity surrounding the amyloid plaque after FGL administration. Similar to the results in hippocampal FGF1 injection, the coverage of LAMP1-immunoreactivity was significantly lower in FGL-injected APP mice than the control-injected APP mice (*p* = 0.0489, Figure 8c, d). Thus, in line with the results from FGF1, FGL has beneficial effects against Aβ induced deficits in multiple cell types.

## 4. Discussion

In this study, we demonstrated that APP mice exhibit lower FGFR1 expression. Activation of FGFR1 with either of the two agonists, FGF1 and FGL, improves AD-related spatial memory deficits, decreases astrogliosis and dystrophic neurites, and increases neurogenesis (Figure 9). Hippocampally-injected FGF1, but not subcutaneously-injected FGL, reduced Aβ deposition *in vivo* and enhanced microglial phagocytosis and Aβ engulfment *in vitro*. These findings indicated that FGFR1 agonists could provide beneficial effects to AD models.

**Figure 9.**
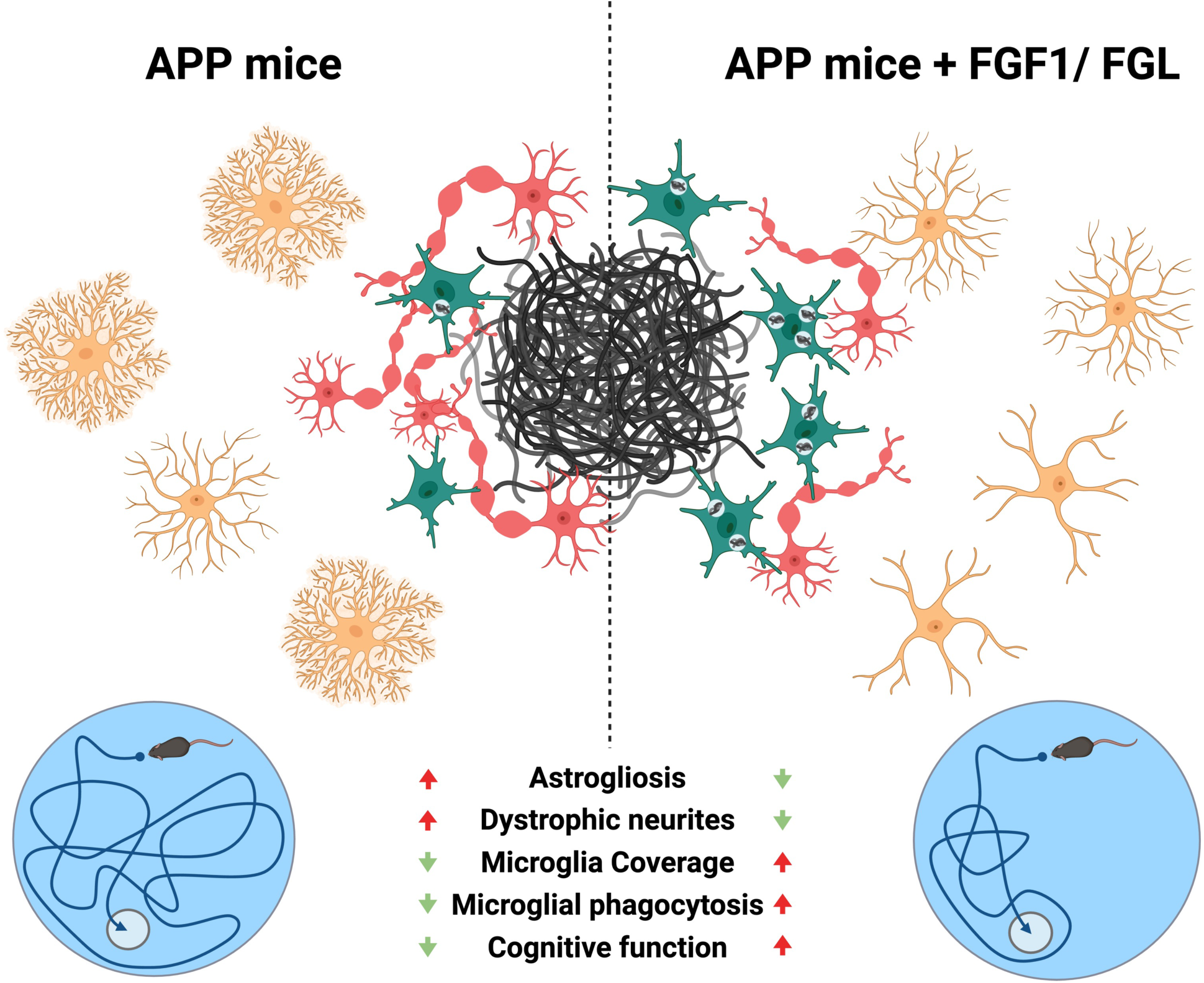
Schematic of FGFR1 agonists mitigates the progression of AD in the APP mouse. In APP mice, accumulation of amyloid plaques triggers astrogliosis, induces dystrophic neurites, and impairs microglia phagocytosis, leading to cognitive impairment. Treatment with FGFR1 agonists effectively reverses these cellular abnormalities and ameliorates memory impairments in APP mice.

FGF1 could interact with FGFR1 (Wu et al., 2003; Zhang et al., 2006), activating downstream signaling pathways that regulate various cellular processes in the brain (Wu et al., 2020). FGL phosphorylates both FGFR1 and FGFR4 *in vitro* (Francavilla et al., 2009). However, given the absence of FGFR4 in the adult brain, the in vivo effects of FGL are likely mediated predominantly through FGFR1 activation. Therefore, our study using FGF1 and FGL as agonists likely predominantly stimulates FGFR1. Expressed in neurons, microglia, and astrocytes, FGFR1 plays a crucial role in regulating the multiple cellular functions in these cell types to control proper brain functions. Our study demonstrated that FGFR1 activation likely improves AD pathology through multiple mechanisms, including promoting neuronal health, reducing astrogliosis, and increasing microglia-related phagocytosis.

Dystrophic neurites, which result from disrupted microtubules and impaired axonal transport due to extracellular plaques, contribute to neuronal dysfunction and death (Adalbert et al., 2009; Sanchez-Varo et al., 2012). The decrease of dystrophic neurites after FGF1 and FGL treatment may be due to the increased formation of the microglia barrier, which could protect the neurites around the plaques. In addition, FGF1 may inhibit the GSK3β-CRMP2 pathway to suppress Aβ-induced neuronal cell death (Cao et al., 2019) and activate the PI3K/Akt signaling pathway to mitigate hypoxic-ischemic neuronal injury (Ye et al., 2019). Furthermore, our findings indicated that FGF1 and FGL can reverse the decline in hippocampal neurogenesis observed in APP mice, consistent with previous studies showing that FGF signaling induces neurogenesis and rescue memory decline in old mice (Kang & Hebert, 2015). Notably, prior research has shown that the FGFR signal promotes neurogenesis through the phospholipase Cγ pathway in response to an enriched environment (Gronska-Peski et al., 2021). However, the precise neuroprotective mechanisms underlying the effects of FGFR agonists in our APP mouse model require further investigation.

Injury-induced astrogliosis could reduce synapse coverage and impair neural connectivity (Sofroniew, 2009). Our study found that FGF1 and FGL reduced astrogliosis in the APP mouse model. The FGF-FGFR signaling is essential for maintaining the resting state of astrocytes in healthy and injured brains (Klimaschewski & Claus, 2021). Deleting all three FGFRs in astrocytes induces glial cell activation and abnormal proliferation (Kang et al., 2014). The potential mechanism of FGFR1 to mitigate astrogliosis is probably mediated through the suppression of the STAT3 pathway (Bi et al., 2024), which reduces astrogliosis and preserves memory functions (Ceyzeriat et al., 2018). These data, in conjunction with our findings, consolidate the role of the FGF-FGFR signaling in regulating astrogliosis.

In AD, dysfunctional microglia fail to clear aggregated Aβ. Our *in vivo* data suggest that FGF1 and FGL treatments increase plaque-associated microglia, which is usually associated with increased phagocytosis (Lin et al., 2024). Echoing the *in vivo* results, our *in vitro* data showed that FGF1 increases the phagocytose ability in both primary microglia and BV2 microglial cell lines. FGF1 has been reported to increase phagocytosis in macrophages (Ichinose et al., 1998a, 1998b). While both macrophages and microglia originate from the myeloid lineage (Chen et al., 2023), this study provides the first demonstration that FGF1 can augment phagocytic activity in microglia. Similarly, FGF2, which also activates FGFR1, enhances microglial migration and phagocytosis of neuronal debris (Kiyota et al., 2011). These data, in conjunction with our findings, indicate that FGFR1 agonists can stimulate phagocytic functions in microglia to clear Aβ.

The primary challenge in developing AD therapeutics is effectively delivering drugs across the BBB using less-invasive approaches. While FGF1 demonstrates promising effects when directly injected into the hippocampus, its large molecular size significantly restricts its ability to cross the BBB following peripheral administration. In contrast, the smaller FGL peptide can cross the BBB rapidly and maintain detectable levels in plasma and CSF for at least 5 hours (Secher et al., 2006). Compared to hippocampal FGF1 administration, subcutaneous FGL injection exhibited beneficial effects on memory deficits but failed to impact Aβ pathology significantly. This discrepancy may be attributed to factors such as the later initiation of FGL treatment compared to FGF1 and the potential suboptimal dosage of FGL administration. Further studies exploring the treatment conditions, such as identifying the ideal treatment window, optimizing the FGL dosage, or adjusting its delivery systems (Meng et al., 2017), are needed to improve intervention outcomes.

## 5. Conclusion

In conclusion, we found that FGFR1 agonists can improve cognitive decline, promote neurogenesis, and reduce dystrophic neurites in AD mouse models regardless of the drug delivery routes. However, only FGF1, when injected directly into the hippocampus, demonstrated effects on reducing amyloid plaques and enhancing phagocytosis. Our study shows that activation of the FGF1-FGFR1 signaling can mitigate AD-related neurodegeneration and foster a more supportive environment for neuronal health. Notably, the small peptide FGL, with its ability to readily cross the BBB, presents a promising therapeutic avenue for AD.

## Acknowledgment

This work was financially supported by the Brain Research Center, National Yang Ming Chiao Tung University from The Featured Areas Research Center Program within the framework of the Higher Education Sprout Project by the Ministry of Education (MOE) in Taiwan; the National Science and Technology Council, Taiwan (NSTC-112-2320-B-A49-015-MY3, MOST 113-2321-B-A49-016; 108-2314-B-532-003; 109-2314-B-532-003), the Departement of Health, Taipei City Government (10901-62-058), and Taipei City Hospital, Ren-I Branch (TPCH-108-16). Behavioral studies were carried out at the Animal Behavioral Core at Brain Research Center, National Yang Ming Chiao Tung University, Taipei, Taiwan. The funders had no role in the study design, data collection and analysis, or manuscript preparation.

## References

2024 Alzheimer’s disease facts and figures. (2024). Alzheimers Dement, 20(5), 3708–3821. 10.1002/alz.13809

Adalbert, R., Nogradi, A., Babetto, E., Janeckova, L., Walker, S. A., Kerschensteiner, M., Misgeld, T., & Coleman, M. P. (2009). Severely dystrophic axons at amyloid plaques remain continuous and connected to viable cell bodies. Brain, 132(Pt 2), 402–416. 10.1093/brain/awn312

Anacker, C., & Hen, R. (2017). Adult hippocampal neurogenesis and cognitive flexibility - linking memory and mood. Nat Rev Neurosci, 18(6), 335–346. 10.1038/nrn.2017.45

Araki, T., Ikegaya, Y., & Koyama, R. (2021). The effects of microglia- and astrocyte-derived factors on neurogenesis in health and disease. Eur J Neurosci, 54(5), 5880–5901. 10.1111/ejn.14969

Bi, J., Wang, Y., Wang, K., Sun, Y., Ye, F., Wang, X., & Pan, J. (2024). FGF1 attenuates sepsis-induced coagulation dysfunction and hepatic injury via IL6/STAT3 pathway inhibition. Biochim Biophys Acta Mol Basis Dis, 1870(7), 167281. 10.1016/j.bbadis.2024.167281

Cambon, K., Hansen, S. M., Venero, C., Herrero, A. I., Skibo, G., Berezin, V., Bock, E., & Sandi, C. (2004). A synthetic neural cell adhesion molecule mimetic peptide promotes synaptogenesis, enhances presynaptic function, and facilitates memory consolidation. J Neurosci, 24(17), 4197–4204. 10.1523/JNEUROSCI.0436-04.2004

Cameron, B., & Landreth, G. E. (2010). Inflammation, microglia, and Alzheimer’s disease. Neurobiol Dis, 37(3), 503–509. 10.1016/j.nbd.2009.10.006

Cao, Q., Meng, T., Man, J., Peng, D., Chen, H., Xiang, Q., Su, Z., Zhang, Q., & Huang, Y. (2019). aFGF Promotes Neurite Growth by Regulating GSK3beta-CRMP2 Signaling Pathway in Cortical Neurons Damaged by Amyloid-beta. J Alzheimers Dis, 72(1), 97–109. 10.3233/JAD-190458

Ceyzeriat, K., Ben Haim, L., Denizot, A., Pommier, D., Matos, M., Guillemaud, O., Palomares, M. A., Abjean, L., Petit, F., Gipchtein, P., Gaillard, M. C., Guillermier, M., Bernier, S., Gaudin, M., Auregan, G., Josephine, C., Dechamps, N., Veran, J., Langlais, V., … Escartin, C. (2018). Modulation of astrocyte reactivity improves functional deficits in mouse models of Alzheimer’s disease. Acta Neuropathol Commun, 6(1), 104. 10.1186/s40478-018-0606-1

Chen, S., Li, J., Meng, S., He, T., Shi, Z., Wang, C., Wang, Y., Cao, H., Huang, Y., Zhang, Y., Gong, Y., & Gao, Y. (2023). Microglia and macrophages in the neuro-glia-vascular unit: From identity to functions. Neurobiol Dis, 179, 106066. 10.1016/j.nbd.2023.106066

Cheng, I. H., Scearce-Levie, K., Legleiter, J., Palop, J. J., Gerstein, H., Bien-Ly, N., Puolivali, J., Lesne, S., Ashe, K. H., Muchowski, P. J., & Mucke, L. (2007). Accelerating amyloid-beta fibrillization reduces oligomer levels and functional deficits in Alzheimer disease mouse models. J Biol Chem, 282(33), 23818–23828. 10.1074/jbc.M701078200

Condello, C., Yuan, P., Schain, A., & Grutzendler, J. (2015). Microglia constitute a barrier that prevents neurotoxic protofibrillar Abeta42 hotspots around plaques. Nat Commun, 6, 6176. 10.1038/ncomms7176

Deng, S. M., Chen, C. J., Lin, H. L., & Cheng, I. H. (2022). The beneficial effect of synbiotics consumption on Alzheimer’s disease mouse model via reducing local and systemic inflammation. IUBMB Life, 74(8), 748–753. 10.1002/iub.2589

Downer, E. J., Cowley, T. R., Lyons, A., Mills, K. H., Berezin, V., Bock, E., & Lynch, M. A. (2010). A novel anti-inflammatory role of NCAM-derived mimetic peptide, FGL. Neurobiol Aging, 31(1), 118–128. 10.1016/j.neurobiolaging.2008.03.017

Francavilla, C., Cattaneo, P., Berezin, V., Bock, E., Ami, D., de Marco, A., Christofori, G., & Cavallaro, U. (2009). The binding of NCAM to FGFR1 induces a specific cellular response mediated by receptor trafficking. J Cell Biol, 187(7), 1101–1116. 10.1083/jcb.200903030

Fruhwurth, S., Zetterberg, H., & Paludan, S. R. (2024). Microglia and amyloid plaque formation in Alzheimer’s disease - Evidence, possible mechanisms, and future challenges. J Neuroimmunol, 390, 578342. 10.1016/j.jneuroim.2024.578342

Gronska-Peski, M., Goncalves, J. T., & Hebert, J. M. (2021). Enriched Environment Promotes Adult Hippocampal Neurogenesis through FGFRs. J Neurosci, 41(13), 2899–2910. 10.1523/JNEUROSCI.2286-20.2021

Han, R. T., Kim, R. D., Molofsky, A. V., & Liddelow, S. A. (2021). Astrocyte-immune cell interactions in physiology and pathology. Immunity, 54(2), 211–224. 10.1016/j.immuni.2021.01.013

Ho Kim, J., Franck, J., Kang, T., Heinsen, H., Ravid, R., Ferrer, I., Hee Cheon, M., Lee, J. Y., Shin Yoo, J., Steinbusch, H. W., Salzet, M., Fournier, I., & Mok Park, Y. (2015). Proteome-wide characterization of signalling interactions in the hippocampal CA4/DG subfield of patients with Alzheimer’s disease. Sci Rep, 5, 11138. 10.1038/srep11138

Huang, Y., Happonen, K. E., Burrola, P. G., O’Connor, C., Hah, N., Huang, L., Nimmerjahn, A., & Lemke, G. (2021). Microglia use TAM receptors to detect and engulf amyloid beta plaques. Nat Immunol, 22(5), 586–594. 10.1038/s41590-021-00913-5

Ichinose, M., Sawada, M., Sasaki, K., & Oomura, Y. (1998a). Effect of acidic fibroblast growth factor (aFGF) on phagocytosis in mouse peritoneal macrophages. Microbiol Immunol, 42(2), 139–142. 10.1111/j.1348-0421.1998.tb02263.x

Ichinose, M., Sawada, M., Sasaki, K., & Oomura, Y. (1998b). Enhancement of phagocytosis in mouse peritoneal macrophages by fragments of acidic fibroblast growth factor (aFGF). Int J Immunopharmacol, 20(4-5), 193–204. 10.1016/s0192-0561(98)00028-9

Kang, W., Balordi, F., Su, N., Chen, L., Fishell, G., & Hebert, J. M. (2014). Astrocyte activation is suppressed in both normal and injured brain by FGF signaling. Proc Natl Acad Sci U S A, 111(29), E2987–2995. 10.1073/pnas.1320401111

Kang, W., & Hebert, J. M. (2015). FGF Signaling Is Necessary for Neurogenesis in Young Mice and Sufficient to Reverse Its Decline in Old Mice. J Neurosci, 35(28), 10217–10223. 10.1523/JNEUROSCI.1469-15.2015

Kiselyov, V. V., Skladchikova, G., Hinsby, A. M., Jensen, P. H., Kulahin, N., Soroka, V., Pedersen, N., Tsetlin, V., Poulsen, F. M., Berezin, V., & Bock, E. (2003). Structural basis for a direct interaction between FGFR1 and NCAM and evidence for a regulatory role of ATP. Structure, 11(6), 691–701. 10.1016/s0969-2126(03)00096-0

Kiyota, T., Ingraham, K. L., Jacobsen, M. T., Xiong, H., & Ikezu, T. (2011). FGF2 gene transfer restores hippocampal functions in mouse models of Alzheimer’s disease and has therapeutic implications for neurocognitive disorders. Proc Natl Acad Sci U S A, 108(49), E1339–1348. 10.1073/pnas.1102349108

Klementiev, B., Novikova, T., Novitskaya, V., Walmod, P. S., Dmytriyeva, O., Pakkenberg, B., Berezin, V., & Bock, E. (2007). A neural cell adhesion molecule-derived peptide reduces neuropathological signs and cognitive impairment induced by Abeta25-35. Neuroscience, 145(1), 209–224. 10.1016/j.neuroscience.2006.11.060

Klimaschewski, L., & Claus, P. (2021). Fibroblast Growth Factor Signalling in the Diseased Nervous System. Mol Neurobiol, 58(8), 3884–3902. 10.1007/s12035-021-02367-0

Knafo, S., Venero, C., Sanchez-Puelles, C., Pereda-Perez, I., Franco, A., Sandi, C., Suarez, L. M., Solis, J. M., Alonso-Nanclares, L., Martin, E. D., Merino-Serrais, P., Borcel, E., Li, S., Chen, Y., Gonzalez-Soriano, J., Berezin, V., Bock, E., Defelipe, J., & Esteban, J. A. (2012). Facilitation of AMPA receptor synaptic delivery as a molecular mechanism for cognitive enhancement. PLoS Biol, 10(2), e1001262. 10.1371/journal.pbio.1001262

Ko, C. C., Tu, T. H., Wu, J. C., Huang, W. C., Tsai, Y. A., Huang, S. F., Huang, H. C., & Cheng, H. (2018). Functional improvement in chronic human spinal cord injury: Four years after acidic fibroblast growth factor. Sci Rep, 8(1), 12691. 10.1038/s41598-018-31083-4

Kumar, A., Fontana, I. C., & Nordberg, A. (2023). Reactive astrogliosis: A friend or foe in the pathogenesis of Alzheimer’s disease. J Neurochem, 164(3), 309–324. 10.1111/jnc.15565

Lie, P. P. Y., & Nixon, R. A. (2019). Lysosome trafficking and signaling in health and neurodegenerative diseases. Neurobiol Dis, 122, 94–105. 10.1016/j.nbd.2018.05.015

Lin, Y. Y., Chang, W. H., Hsieh, S. L., & Cheng, I. H. (2024). The deficient CLEC5A ameliorates the behavioral and pathological deficits via the microglial Abeta clearance in Alzheimer’s disease mouse model. J Neuroinflammation, 21(1), 273. 10.1186/s12974-024-03253-x

Liu, Y. L., Chen, W. T., Lin, Y. Y., Lu, P. H., Hsieh, S. L., & Cheng, I. H. (2017). Amelioration of amyloid-beta-induced deficits by DcR3 in an Alzheimer’s disease model. Mol Neurodegener, 12(1), 30. 10.1186/s13024-017-0173-0

Mabrouk, R., Miettinen, P. O., & Tanila, H. (2023). Most dystrophic neurites in the common 5xFAD Alzheimer mouse model originate from axon terminals. Neurobiol Dis, 182, 106150. 10.1016/j.nbd.2023.106150

Meng, T., Cao, Q., Lei, P., Bush, A. I., Xiang, Q., Su, Z., He, X., Rogers, J. T., Chiu, I. M., Zhang, Q., & Huang, Y. (2017). Tat-haFGF(14-154) Upregulates ADAM10 to Attenuate the Alzheimer Phenotype of APP/PS1 Mice through the PI3K-CREB-IRE1alpha/XBP1 Pathway. Mol Ther Nucleic Acids, 7, 439–452. 10.1016/j.omtn.2017.05.004

Moon, M., Cha, M. Y., & Mook-Jung, I. (2014). Impaired hippocampal neurogenesis and its enhancement with ghrelin in 5XFAD mice. J Alzheimers Dis, 41(1), 233–241. 10.3233/JAD-132417

Neiiendam, J. L., Kohler, L. B., Christensen, C., Li, S., Pedersen, M. V., Ditlevsen, D. K., Kornum, M. K., Kiselyov, V. V., Berezin, V., & Bock, E. (2004). An NCAM-derived FGF-receptor agonist, the FGL-peptide, induces neurite outgrowth and neuronal survival in primary rat neurons. J Neurochem, 91(4), 920–935. 10.1111/j.1471-4159.2004.02779.x

Pankratova, S., Bjornsdottir, H., Christensen, C., Zhang, L., Li, S., Dmytriyeva, O., Bock, E., & Berezin, V. (2016). Immunomodulator CD200 Promotes Neurotrophic Activity by Interacting with and Activating the Fibroblast Growth Factor Receptor. Mol Neurobiol, 53(1), 584–594. 10.1007/s12035-014-9037-6

Patani, R., Hardingham, G. E., & Liddelow, S. A. (2023). Functional roles of reactive astrocytes in neuroinflammation and neurodegeneration. Nat Rev Neurol, 19(7), 395–409. 10.1038/s41582-023-00822-1

Pereda-Perez, I., Valencia, A., Baliyan, S., Nunez, A., Sanz-Garcia, A., Zamora, B., Rodriguez-Fernandez, R., Esteban, J. A., & Venero, C. (2019). Systemic administration of a fibroblast growth factor receptor 1 agonist rescues the cognitive deficit in aged socially isolated rats. Neurobiol Aging, 78, 155–165. 10.1016/j.neurobiolaging.2019.02.011

Podlesny-Drabiniok, A., Marcora, E., & Goate, A. M. (2020). Microglial Phagocytosis: A Disease-Associated Process Emerging from Alzheimer’s Disease Genetics. Trends Neurosci, 43(12), 965–979. 10.1016/j.tins.2020.10.002

Popov, V. I., Medvedev, N. I., Kraev, I. V., Gabbott, P. L., Davies, H. A., Lynch, M., Cowley, T. R., Berezin, V., Bock, E., & Stewart, M. G. (2008). A cell adhesion molecule mimetic, FGL peptide, induces alterations in synapse and dendritic spine structure in the dentate gyrus of aged rats: a three-dimensional ultrastructural study. Eur J Neurosci, 27(2), 301–314. 10.1111/j.1460-9568.2007.06004.x

Saffell, J. L., Williams, E. J., Mason, I. J., Walsh, F. S., & Doherty, P. (1997). Expression of a dominant negative FGF receptor inhibits axonal growth and FGF receptor phosphorylation stimulated by CAMs. Neuron, 18(2), 231–242. 10.1016/s0896-6273(00)80264-0

Salta, E., Lazarov, O., Fitzsimons, C. P., Tanzi, R., Lucassen, P. J., & Choi, S. H. (2023). Adult hippocampal neurogenesis in Alzheimer’s disease: A roadmap to clinical relevance. Cell Stem Cell, 30(2), 120–136. 10.1016/j.stem.2023.01.002

Sanchez-Varo, R., Trujillo-Estrada, L., Sanchez-Mejias, E., Torres, M., Baglietto-Vargas, D., Moreno-Gonzalez, I., De Castro, V., Jimenez, S., Ruano, D., Vizuete, M., Davila, J. C., Garcia-Verdugo, J. M., Jimenez, A. J., Vitorica, J., & Gutierrez, A. (2012). Abnormal accumulation of autophagic vesicles correlates with axonal and synaptic pathology in young Alzheimer’s mice hippocampus. Acta Neuropathol, 123(1), 53–70. 10.1007/s00401-011-0896-x

Secher, T., Novitskaia, V., Berezin, V., Bock, E., Glenthøj, B., & Klementiev, B. (2006). A neural cell adhesion molecule-derived fibroblast growth factor receptor agonist, the FGL-peptide, promotes early postnatal sensorimotor development and enhances social memory retention. Neuroscience, 141(3), 1289–1299. 10.1016/j.neuroscience.2006.04.059

Simpson, J. E., Ince, P. G., Lace, G., Forster, G., Shaw, P. J., Matthews, F., Savva, G., Brayne, C., Wharton, S. B., Function, M. R. C. C., & Ageing Neuropathology Study, G. (2010). Astrocyte phenotype in relation to Alzheimer-type pathology in the ageing brain. Neurobiol Aging, 31(4), 578–590. 10.1016/j.neurobiolaging.2008.05.015

Smit, T., Deshayes, N. A. C., Borchelt, D. R., Kamphuis, W., Middeldorp, J., & Hol, E. M. (2021). Reactive astrocytes as treatment targets in Alzheimer’s disease-Systematic review of studies using the APPswePS1dE9 mouse model. Glia, 69(8), 1852–1881. 10.1002/glia.23981

Sofroniew, M. V. (2009). Molecular dissection of reactive astrogliosis and glial scar formation. Trends Neurosci, 32(12), 638–647. 10.1016/j.tins.2009.08.002

Wu, J. C., Huang, W. C., Chen, Y. C., Tu, T. H., Tsai, Y. A., Huang, S. F., Huang, H. C., & Cheng, H. (2011). Acidic fibroblast growth factor for repair of human spinal cord injury: a clinical trial. J Neurosurg Spine, 15(3), 216–227. 10.3171/2011.4.SPINE10404

Wu, Y., Wu, C., Ye, L., Wang, B., Yuan, Y., Liu, Y., Zheng, P., Xiong, J., Li, Y., Jiang, T., Li, X., & Xiao, J. (2020). Exogenous fibroblast growth factor 1 ameliorates diabetes-induced cognitive decline via coordinately regulating PI3K/AKT signaling and PERK signaling. Cell Commun Signal, 18(1), 81. 10.1186/s12964-020-00588-9

Wu, Z. L., Zhang, L., Yabe, T., Kuberan, B., Beeler, D. L., Love, A., & Rosenberg, R. D. (2003). The involvement of heparan sulfate (HS) in FGF1/HS/FGFR1 signaling complex. J Biol Chem, 278(19), 17121–17129. 10.1074/jbc.M212590200

Xie, Y., Su, N., Yang, J., Tan, Q., Huang, S., Jin, M., Ni, Z., Zhang, B., Zhang, D., Luo, F., Chen, H., Sun, X., Feng, J. Q., Qi, H., & Chen, L. (2020). FGF/FGFR signaling in health and disease. Signal Transduct Target Ther, 5(1), 181. 10.1038/s41392-020-00222-7

Ye, L., Wang, X., Cai, C., Zeng, S., Bai, J., Guo, K., Fang, M., Hu, J., Liu, H., Zhu, L., Liu, F., Wang, D., Hu, Y., Pan, S., Li, X., Lin, L., & Lin, Z. (2019). FGF21 promotes functional recovery after hypoxic-ischemic brain injury in neonatal rats by activating the PI3K/Akt signaling pathway via FGFR1/beta-klotho. Exp Neurol, 317, 34–50. 10.1016/j.expneurol.2019.02.013

Zakrzewska, M., Wiedlocha, A., Szlachcic, A., Krowarsch, D., Otlewski, J., & Olsnes, S. (2009). Increased protein stability of FGF1 can compensate for its reduced affinity for heparin. J Biol Chem, 284(37), 25388–25403. 10.1074/jbc.M109.001289

Zhang, X., Ibrahimi, O. A., Olsen, S. K., Umemori, H., Mohammadi, M., & Ornitz, D. M. (2006). Receptor specificity of the fibroblast growth factor family. The complete mammalian FGF family. J Biol Chem, 281(23), 15694–15700. 10.1074/jbc.M601252200

Zhao, M., Li, D., Shimazu, K., Zhou, Y. X., Lu, B., & Deng, C. X. (2007). Fibroblast growth factor receptor-1 is required for long-term potentiation, memory consolidation, and neurogenesis. Biol Psychiatry, 62(5), 381–390. 10.1016/j.biopsych.2006.10.019

Zhou, H., Pan, H., Li, X., Huang, L., Zhang, R., Yan, X., & Xu, J. (2025). Ginsenoside reprogramming microglia through the FGF/FGFR1 inhibits post traumatic stress disorder. Int Immunopharmacol, 145, 113763. 10.1016/j.intimp.2024.113763

Zou, L. H., Shi, Y. J., He, H., Jiang, S. M., Huo, F. F., Wang, X. M., Wu, F., & Ma, L. (2019). Effects of FGF2/FGFR1 Pathway on Expression of A1 Astrocytes After Infrasound Exposure. Front Neurosci, 13, 429. 10.3389/fnins.2019.00429

